# Thick Filament Activation Is Different in Fast and Slow-Twitch Skeletal Muscle

**DOI:** 10.1101/2022.07.09.499423

**Authors:** Henry M. Gong, Weikang Ma, Michael Regnier, Thomas C. Irving

## Abstract

The contractile properties of fast-twitch and slow-twitch skeletal muscles are primarily determined by the myosin isoform content and modulated by a variety of sarcomere proteins. X-ray diffraction studies of regulatory mechanisms in muscle contraction have focused predominately on fast-or mixed-fiber muscle with slow muscle being much less studied. Here, we used time-resolved x-ray diffraction to investigate the dynamic behavior of the myofilament proteins in relatively pure slow fiber rat soleus (SOL) and pure fast fiber rat extensor digitorum longus (EDL) muscle during twitch and tetanic contractions at optimal length. During twitch contractions the diffraction signatures indicating a transition in the myosin heads from ordered OFF states, where heads are held close to the thick filament backbone, to disordered ON states, where heads are free to bind to thin filaments, were found in EDL and not in SOL muscle. During tetanic contraction, changes in the disposition of myosin heads as active tension develops is a cooperative process in EDL muscle whereas in SOL muscle this relationship is less cooperative. The observed reduced extensibility of the thick filaments in SOL muscle as compared to EDL muscles indicate a molecular basis for this behavior. These data indicate that for the EDL thick filament activation is a cooperative strain-induced mechano-sensing mechanism, whereas for the SOL thick filament activation has a more graded response. These different approaches to thick filament regulation in fast- and slow-twitch muscles may be adaptations for short duration, strong contractions versus sustained, finely controlled contractions, respectively.

**Key Points:**

- Fast twitch muscle and slow-twitch muscle are optimized for strong, short duration contractions and sustained smaller movements respectively.
- Structural events (OFF to ON transitions) in the myosin containing thick filaments in fast muscle help determine the timing and strength of contractions but these have not been studied in slow-twitch muscle.
- The X-ray diffraction signatures of structural OFF to ON transitions are different in fast extensor digitorum longus (EDL) and slow soleus (SOL) muscle, being completely absent during twitches in soleus muscle and blunted during tetanic contractions SOL as compared to EDL
- Cooperative thick filament structural OFF to ON transitions in fast twitch muscle may be an adaptation for rapid and ballistic movements whereas more graded OFF to ON structural transitions in slow-twitch muscle may be an adaptation for slower, finer motions.

## Introduction

Contraction in striated muscles is the result of myosin motor molecules on the thick filaments interacting with the binding sites on the actin-containing thin filaments in the sarcomere (Gordon *et al*., 2000). Muscle contraction is initiated when intracellular Ca^2+^ released from the sarcoplasmic reticulum bind to troponin C. This leads to a series of interactions within the troponin complex that results in increased mobility of tropomyosin on the F-actin strands of thin filaments. This, in turn, results in increased exposure of myosin binding sites on F-actin for the formation and cycling of cross-bridges (Gordon *et al*., 2000). However, muscle regulation is now understood to require both thick and thin filament activation mechanisms. Under resting conditions, myosin heads are arranged in a quasi-helical array around the thick filament backbone, an OFF state unable to interact with actin (Huxley & Brown, 1967). The calcium-mediated thin filament-based regulatory mechanism, however, does not address the question of how the myosin heads in the OFF state, are released to an ON state to interact with actin. It is known that thick filament activation mechanisms that trigger the conversion of myosin heads from the OFF to the ON state vary depending on the muscle types. For example, in some invertebrates such as scallop, myosin can be activated by direct binding of calcium to the regulatory light chains (RLC) (Szent-Györgyi, 2007). Tarantula leg muscle, another invertebrate muscle, activates its thick filament through binding of calcium to calmodulin and thereby activating myosin light chain kinase (MLCK) that phosphorylated the myosin RLC (Padrón *et al*., 2020).

In contrast to invertebrate muscles, vertebrate skeletal and cardiac muscles have been proposed to utilize a mechano-sensing mechanism (Linari *et al*., 2015; Irving, 2017). In a resting muscle, a large percentage of myosin heads are in the structurally defined OFF state, a quasi-helically ordered state in which myosin molecules on the surface of the thick filament give rise to myosin-based layer line reflections. This helical ordering is lost when myosin heads are turned ON to participate in contraction (Huxley & Brown, 1967; Linari et al., 2015; Ma et al., 2018a). The mechano-sensing model proposes that once thin filaments are turned ON, a small number of constitutively ON myosin heads generate a strain on the thick filament backbone, evidence by the increases in the spacings of the M6 meridional reflections. It is this strain that is proposed to release the myosin heads from the OFF state, as evidenced by the decrease in the intensity of the first myosin layer lines (IMLL1) and an increase in the spacing of the M3 meridional. These ON-state myosin heads are now free to bind to actin and generate force (Linari *et al*., 2015).

Skeletal muscles in mammals consist of a mixture of fast- and slow-twitch fibers, which differ in architecture and metabolic properties. For example, fast-twitch fibers can generate high amounts of force for short periods of time (Rome *et al*., 1988; Schiaffino & Reggiani, 2011) in ballistic movements, while slow-twitch fibers usually contract in a more graded way and are more resistant to fatigue (Schiaffino & Reggiani, 2011). Most structural studies of vertebrate muscle have focused on muscles with predominantly fast-twitch fibers, or those with a mixture of fast-twitch and slow-twitch fibers (Gutmann, 1966; Salviati *et al*., 1982; Hämäläinen & Pette, 1993; Lutz *et al*., 1998; Linari *et al*., 2015). In contrast, the structural events during contraction in slow-twitch fibers have been much less studied.

We compared the sequence of structural events in the sarcomeres of extensor digitorum longus (EDL) and soleus (SOL) muscle fibers from the hindlimbs of adult rats at rest and during twitch, and tetanic contractions using time-resolved x-ray diffraction. SOL muscle is comprised of 85-95% slow type-1 fibers and 5-15% fast-2A fibers (Wigston & English, 1992; Li *et al*., 2019). The EDL is made up of 100% type-2 fast-twitch fibers (Eng *et al*., 2008; Li *et al*., 2019). The relatively pure composition of fiber types in rat EDL and SOL makes them an appropriate model to study fiber-type differences in thick filament activation. Here, we report that the sequence of molecular events during twitches and tetanic contraction differ in fast EDL muscle and slow SOL. Thick filament activation in EDL is consistent with the mechano-sensing model in both twitches and tetanic contractions. During twitches in SOL muscle, there are few indications of thick filament activation with the force most likely generated by constitutively ON heads. During tetani, thick filaments become more activated but this process is much less cooperative, consistent with a graded continuous response rather than a stepwise transition at a certain value of thick filament strain. These results indicate that thick filament regulation is not the same in all muscle types, and this most likely reflects optimization for their physiological function.

## Materials and Methods

### Muscle preparation and experimental apparatus

All animal experiments were done in compliance with protocols approved by the Illinois Institute of Technology Institutional Animal Care and Use Committees and followed the “Guide for the Care and Use of Laboratory Animals” published by the US National Institutes of Health (NIH Publication No. 85-23, revised 2011) Rats were euthanized and the muscles were isolated as described previously (Ma & Irving, 2019). EDL and SOL muscles were isolated from the hindlimbs and split into 3-4 smaller bundles. Intact leg muscles in rats are covered with collagenous fascia. By splitting the whole muscle into smaller bundles, these collagenous regions can be largely avoided, improving the quality of the x-ray patterns. Fiber bundles were tied with sutures at both ends and transferred to a chamber containing Ringer’s solution (145 mM NaCl, 2.5 mM KCl, 0.9 mM CaCl_2_H_2_O, 1 mM MgSO_4_7H_2_O, 10.9 mM Glucose, 9.9 mM HEPES, pH 7.4). The suture attached to one end of the muscle was tied to a fixed hook in the chamber. The other end of the muscle bundle was attached via its suture to a servo motor/force transducer (300C, Aurora Scientific Inc, Aurora, Canada). The chamber was continuously bubbled with oxygen and the platinum electrodes attached to a biphasic current stimulator (model 710A, Aurora Scientific Inc). Length was controlled, and force recorded, using an ASI 610A data acquisition and control system (Aurora Scientific Inc). All experiments were performed at room temperature (23°C).

### X-ray diffraction

X-ray diffraction experiments were performed using the small angle instrument on the BioCAT Beamline 18ID at the Advanced Photon Source, Argonne National Laboratory, Lemont, Illinois (Fischetti *et al*., 2004). The x-ray beam energy was set at 12 keV (0.1033 nm wavelength) with a flux of ∼10^13^ photons per second. The x-ray beam was focused to 250 × 250 µm at the sample position and 30 × 120 µm at the detector position. The diffraction patterns were collected in two separate experimental runs. The first experiment used a sample-to-detector distance of 6 m. The other experiment had a sample-to-detector distance of 3.6 m. All x-ray patterns were collected with a Pilatus 3 1M detector (Dectris, Baden-Daettwil, Switzerland).

The stimulator was adjusted to provide an optimal current and voltage for each sample as judged by the settings where the sample generated maximum twitch force. Then the sample was stretched to its optimal length (L_o_), defined as the length where maximum twitch force was generated. EDL muscles were stimulated after 0.05 s of initial delay with a pulse width of 0.2 ms. SOL muscles were stimulated after a 0.5 s of initial delay with a pulse width of 0.2 ms. X-ray patterns from EDL and SOL muscles were collected continuously with a 0.008 s exposure time and 0.01 s exposure period for a frame rate of 100 Hz. In order to minimize radiation damage, the samples were oscillated during x-ray exposure at 10 mm/s. After each experiment, the muscle was fixed using 10% formalin at its L_o_ for sarcomere length (SL) measurement. SL was measured by extracting a small bundle of fibers from the fixed muscle bundle and observing it under an inverted microscope with a 40X objective and the bright field striation pattern analyzed with a High-Speed Video Sarcomere Length (HVSL) system (Aurora Scientific Inc).

### Data analysis

Calibrated force values were extracted from raw data traces using ASI 611A Dynamic Muscle Analysis v5.300 software (Aurora Scientific Inc). The cross-sectional area of the muscle was calculated as described previously with the assumption that muscle density is 1.056 mg/mm^2^ (Ward & Lieber, 2005; Eng *et al*., 2008). Tension was calculated by dividing the force developed by the cross-section area of the preparation. The rate of the tension rise, and tension decay were obtained by manually finding the half-time (1/2 time) after normalizing the smallest value as 0 and biggest value as 100%. For twitch contractions, the data points between resting-tension (RE) and peak-tension (PT) were compared. The ½ times of the twitch contractions were obtained by fitting to one-phase decay. In tetanic contractions, the data values between RE and the plateau region of tension (PLA) were compared. The ½ times of the tetanic contractions were obtained by fitting to a Hill function. All graphs were plotted with mean ± SEM and ½ times were reported with 95% confidence intervals (CI). All statistical analyses were performed using GraphPad Prism 9 (GraphPad Software). All the data was analyzed using Student’s t-tests. Symbols on the figures: ns: p ≥ 0.05, *: p < 0.05, **: p < 0.01, and ***: p < 0.001, and ***: p < 0.0001.

X-ray diffraction patterns were analyzed using the MuscleX software suite developed by BioCAT (Jiratrakanvong *et al*., 2018). The equatorial reflections were analyzed using the Equator routine in the MuscleX suite. The spacings of meridional reflections were analyzed using the Projection Traces routine in the MuscleX suite. The intensity of first myosin-based layer line (MLL1) and the first actin-based layer line (ALL1) overlap with each other in the diffraction. To measure their intensity separately, the projected intensity traced parallel to the meridian was fitted to a double-Gaussian curve under the peak using FITYK (Wojdyr, 2010). The sixth actin-based layer lines (ALL6) and seventh actin-based layer line (ALL7) were analyzed as described previously (Wakabayashi et al. 1994). The axial spacing of the actin subunits was calculated based on the change in ALL6 spacing and ALL7 spacing between resting and contraction as previously been described (Eq.1) (Egelman *et al*., 1982). All spacing values, and intensity values are given as means ± SEM. The rate values are given means and its corresponding confidence interval (95% CI).

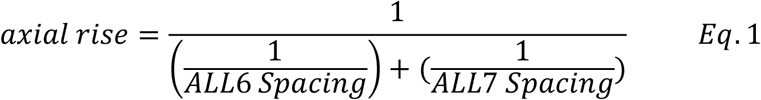

## Results

### X-ray diffraction patterns from EDL and SOL muscle

The x-ray diffraction patterns of EDL (Figure 1A and C, left panel) and SOL muscles (Figure 1B and D, left panel) show prominent reflections along the meridional axis and the equatorial axis, as well as myosin-based layer lines parallel to the equator under resting conditions. The layer lines from SOL muscle have pronounced lattice sampling where the intensity maxima on the layer lines are aligned with 1,0 and 1,1 reflections on the equator. In contrast, the diffraction maxima on the layer lines from EDL muscle do not line up with the equatorial reflections. These observations agree with a previous report that EDL muscle has a super-lattice, and SOL has a simple lattice (Ma *et al*., 2019). When activated, the layer lines during twitch contraction remain prominent for both muscles (Figure 1A and B, right panel), whereas in tetanic contraction the intensity of the layer lines decrease substantially but still remain visible (Figure 1C and D, right panel) (Ma *et al*., 2018*a*). The third myosin-based meridional reflection (M3), which arises from the axial separation of myosin heads along the thick filament, and the sixth myosin-based meridional reflection (M6), which arises from structures within the backbone of the thick filament, moved inward towards the center of the pattern during contraction (Linari *et al*., 2000; Huxley, 2004; Reconditi, 2006). This spacing increase is more pronounced during tetanic contraction as compared to twitch, and indicates an increase in the axial periodicity in the thick filament.

**Figure 1.**
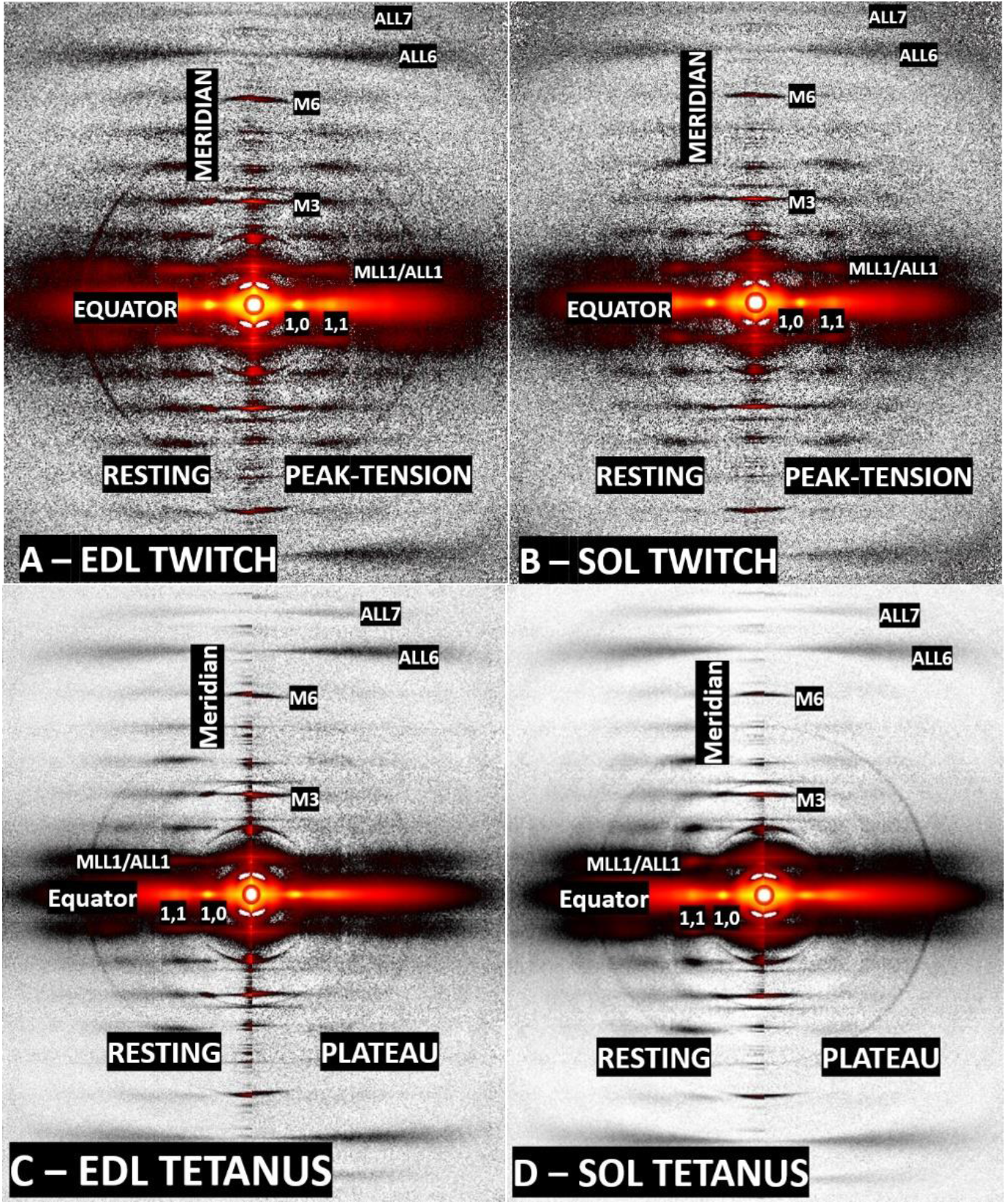
X-ray diffraction patterns during twitch and tetanic contraction. The top two images are the x-ray patterns collected from EDL (A) and SOL (B) during twitch contraction. Each image consists of patterns from resting (left half) and peak-tension (right half). The bottom two images are patterns from EDL (C) and SOL (D) collected during tetanic contraction. Same as the top two images, the left half is from resting and the right half is from the plateau region.

### Structural change during fixed-end twitch contractions

At L_o_, the resting sarcomere lengths were 2.73 ± 0.03 μm and 2.81 ± 0.02 μm for EDL and SOL respectively. These sarcomere lengths at L_o_ are similar to values reported previously (Close, 1964, 1972). Resting tension at L_0_ was significantly greater for SOL compared with EDL (p < 0.0001). The higher passive tension at rest observed in SOL is because the muscle was required to be stretched to a longer sarcomere length to reach Lo as previously reported for mouse SOL muscle (Kiss *et al*., 2018). During twitch contractions (**Error! Reference source not found**.), EDL muscle (**Error! Reference source not found**.A) generated an active tension of 19.4 ± 2.72 mN/mm^2^. Tension developed with a half-time of 16.1 ms (95% CI: 14.9 ms to 17.3 ms) and decayed with a half-time of 52.4 ms (95% CI: 46.2 ms to 58.7 ms). SOL muscles (Figure 3A) produced an active tension of 13 ± 3.29 mN/mm^2^ with a half-time of 29.7 ms (95% CI: 27.4 ms to 32 ms). After tension peaked, it decayed with a half-time of 659.4 ms (95% CI: 502.8 ms to 815.9 ms). The twitch tensions for both muscles reported here are comparable with those reported previously (Hallal *et al*., 2019).

The hexagonal crystal lattice formed by the myosin-containing thick and actin-containing thin filaments in the sarcomere give rise to the two strongest equatorial reflections, the 1,0 and the 1,1. The intensity ratio (I_1,1_/I_1,0_) has been demonstrated to closely track active force in many striated muscles (Matsubara, 1980; Brenner & Yu, 1985; Ma *et al*., 2018*a*), therefore it is often used as a measure of the degree of association of myosin heads with actin during contraction. The I_1,1_/I_1,0_ of EDL increased from 0.38 ± 0.02 to 0.51 ± 0.03 (p < 0.0001, Figure 2B) at peak-tension and 0.38 ± 0.03 to 0.46 ± 0.03 (p = 0.0006, Figure 3B) for SOL. Due to insufficient time resolution between detector frames (10 ms/frame), the rate of the ascending portion cannot be reliable obtained for both EDL and SOL, but the clear increases in I_1,1_/I_1,0_ with tension indicate that the myosin heads have left the vicinity of the thick filament backbones and are in the vicinity of thin filaments during twitch contraction. After reaching peak-tension, I_1,1_/I_1,0_ decays with a half-time of 58.9 ms (95% CI: 33.9 ms to 155.8 ms) and 560.2 ms (95% CI: 391 ms to 897.1 ms) for EDL muscle SOL muscle respectively. The decay of I_1,1_/I_1,0_ for EDL muscle closely tracks tension decay, whereas for soleus I_1,1_/I_1,0_ decays faster than tension.

**Figure 2.**
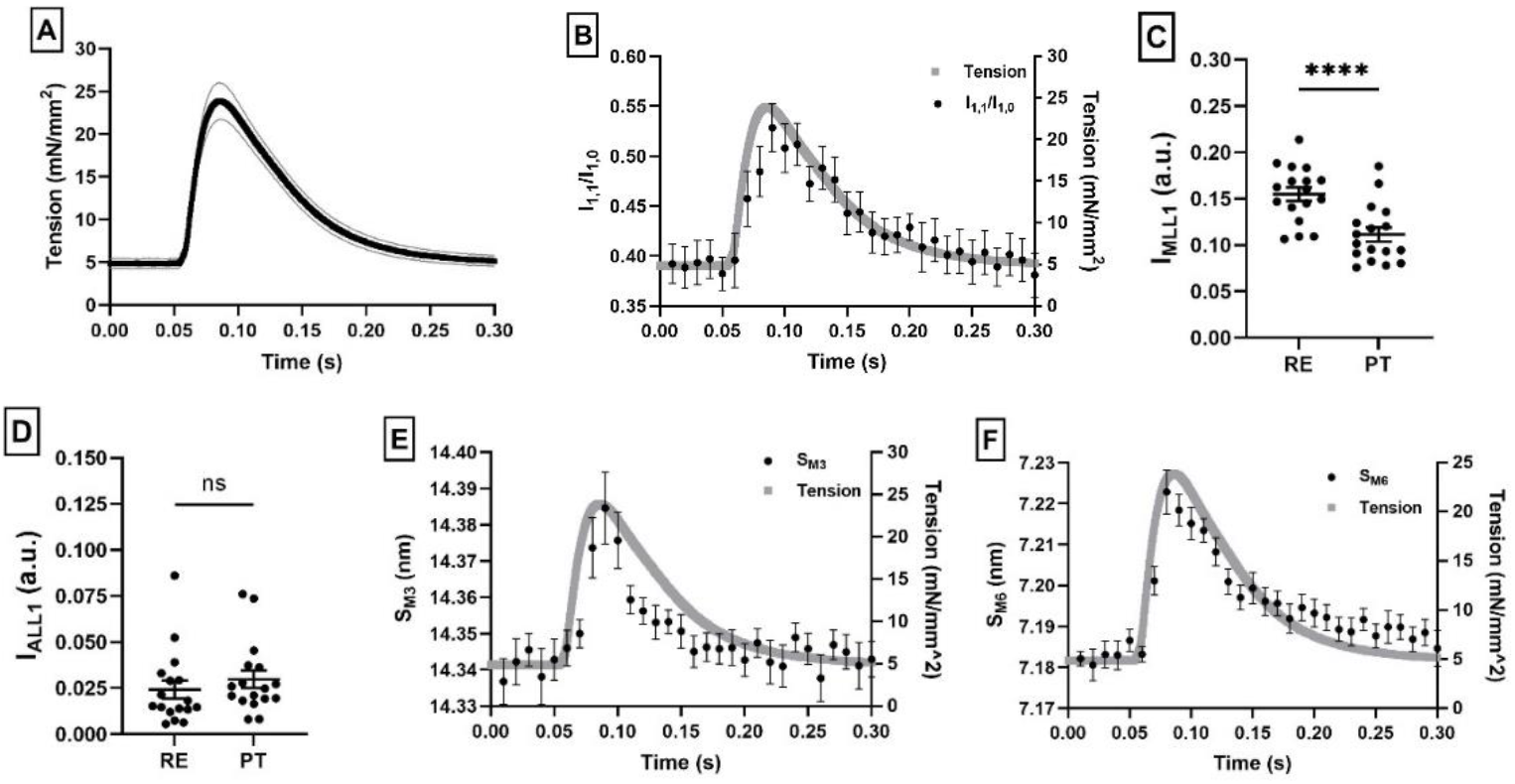
X-ray reflections of EDL during twitch activation. As tension (A) develops, the intensity ratio (B), the spacing of M3 (S_M3_, E), and the spacing of M6 (S_M6_, F) all increase. The corresponding MLL1 intensity (I_MLL1_, C) decreases and ALL1 intensity (I_ALL1_, D) increases. All graphs are plotted with mean ± SEM, n = 9.

**Figure 3.**
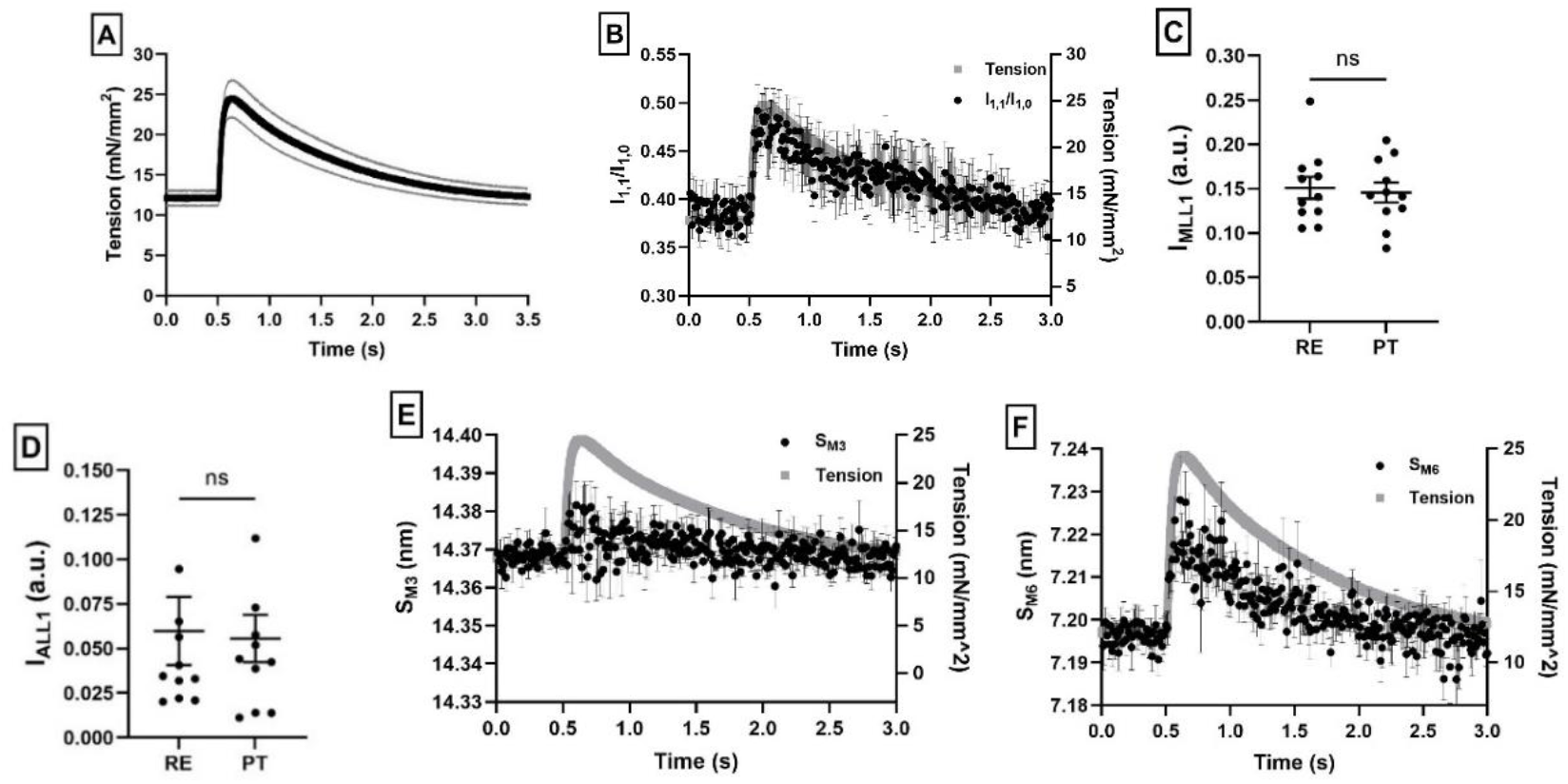
X-ray reflections of SOL during twitch activation. As tension (A) develops, the intensity ratio (B), and the spacing of M6 (S_M6_, F) all increase. The corresponding MLL1 intensity (I_MLL1_, C), ALL1 intensity (I_ALL1_, D), and the spacing of M3 (S_M3_, E) remain the same. All data are plotted as mean ± SEM, n = 9.

The first myosin-based layer-line (MLL1) is one of the prominent diffraction features that is closest to the center of the pattern (Figure 1). It originates primarily from the quasi-helical arrangement of the myosin heads on the surface of the thick filament with a periodicity of ∼42.9 nm (Huxley & Brown, 1967; Caremani *et al*., 2019; Ma *et al*., 2020). The first actin-based layer-line (ALL1) arises from the thin filament double helical structure with a helical repeat of ∼35.9 nm (Dominguez & Holmes, 2011). When muscle contracts, the intensity of MLL1 (I_MLL1_) decreases and the intensity of ALL1 (I_ALL1_) increases (Wakabayashi *et al*., 1988). This decrease in I_MLL1_ reflects a loss of helical ordering due to the departure of myosin heads from their positions near the thick filament backbone. The increase in I_ALL1_ results from the formation of cross-bridges (Tsaturyan *et al*., 2011). In EDL, I_MLL1_ decreased significantly from 0.15 ± 0.01 at rest to 0.11 ± 0.01 at peak-tension (p < 0.0001, Figure 2C). I_ALL1_ appeared to increase during to contraction, but this was not significant (from 0.024 ± 0.005 to 0.03 ± 0.005 (p = 0.1443, Figure 2D). Similar observations have been reported in mouse muscle (Hill *et al*., 2021). In contrast to EDL muscle, I_MLL1_ in SOL muscle did not change significantly (0.15 ± 0.01 to 0.146 ± 0.01, p = 0.5870, Figure 3C). I_ALL1_ in SOL muscle also did not change significantly _(_0.06 ± 0.02 to 0.056 ± 0.01, p = 0.7437, Figure 3D). These results indicate that the helical ordering of myosin heads on the thick filament was preserved during fixed end twitch contractions in SOL whereas there is a substantial reduction of head ordering in EDL.

The quasi-helical arrangement of the three-stranded myosin heads on the surface of the thick filament also produce x-ray reflections on the meridian, the most prominent of which is the third order myosin meridional reflection (M3, Figure 1). It arises primarily from the ∼14.3 nm axial separation of myosin heads and is often used to extract structural information concerning the myosin heads during force development (e.g. (Linari *et al*., 2000; Huxley, 2004; Reconditi, 2006). During twitch contraction, S_M3_ from EDL increased from 14.34 ± 0.01 nm at rest to 14.38 ± 0.01 nm at peak tension (p < 0.0003, Figure 2E), whereas there was no significant increase of S_M3_ in SOL from rest to peak-tension (14.368 ± 0.002 nm to 14.374 ± 0.003 nm, p = 0.0694, Figure 3E) (Honda *et al*., 1996). Due to insufficient time resolution, the rate of change of S_M3_ during twitch contraction cannot be reliably estimated but the significant increase in S_M3_ provides clear evidence that the myosin heads in EDL have moved towards an active ON configuration (Hill *et al*., 2021). In the case of SOL muscle, the lack of an obvious increase in S_M3_ with tension development suggest that the majority of the myosin heads have retained their resting periodicity (∼14.3 nm) throughout the twitch contraction. This agrees with previous findings from skinned rat SOL muscle (Honda *et al*., 1996). After achieving peak-tension, S_M3_ of EDL decays with a half-time of 18.9 ms (95% CI: 11.9 ms to 30.7 ms).

The sixth order meridional reflection (M6) is another prominent feature on the meridian that arises primarily from structures within the thick filament backbone with a periodicity of ∼7.2 nm (Linari *et al*., 2000; Huxley, 2004; Reconditi, 2006). The spacing change of M6 is interpreted as the extensibility of backbone as a function of strain in the thick filament generated by either passive or active force (Huxley *et al*., 1994; Wakabayashi *et al*., 1994; Linari *et al*., 2000; Huxley, 2004; Ma *et al*., 2018*b*). An increase in S_M6_ is also associated with activation of the thick filament (Haselgrove, 1975). During twitch contractions in EDL, S_M6_ increased from 7.19 ± 0.001 nm at rest to 7.22 ± 0.004 nm at peak-tension (p < 0.0001, Figure 2F). The increase in S_M6_ in SOL was smaller as compared to EDL, changing from 7.195 ± 0.002 nm at rest to 7.214 ± 0.004 nm at peak-tension (P = 0.0001, Figure 3F). After peak-tension, S_M6_ decays with a half-time of 43.9 ms (95% CI: 31.3 ms to 67.1 ms) and 624.1 ms (95% CI: 470.3 ms to 891.2 ms) for EDL and SOL muscle, respectively.

### Structural changes during fixed-end tetanic contractions

In the twitch experiments it appeared that the changes in diffraction features associated with the mechano-sensing model were observed in EDL muscle but not in SOL. To further investigate this phenomenon, and whether these results would be similar with more vigorous contractions, we performed tetanic contraction experiments, where SOL and EDL were tetanically activated. At maximum tetanic tension, EDL muscle generated 84.43 ± 8.94 mN/mm^2^ (Table 1, Figure 4A). The half-time of tension development was 45 ms (95% CI: 42.1 ms to 47.9 ms) and the half-time of relaxation was 57.6 ms (95% CI: 55 ms to 60.2 ms). For SOL muscle, maximum active tension was 93.36 ± 9.57 mN/mm^2^ (Figure 5A). The half-times were 197.2 ms (95% CI: 165.6 ms to 228.9 ms) and 217.4 ms (95% CI: 183.1 ms to 251.7 ms) for tension development and relaxation, respectively. Force development and relaxation is faster in fast-twitch EDL than in slow-twitch SOL muscle in agreement with results reported previously (Wallinga-de Jonge *et al*., 1980).

**Table 1.** Tension values extracted during resting and contraction. The top half of this table contains the tension values from twitch contraction. The bottom half are the values from tetanic contraction. The active tension is calculated as difference between contraction and resting. All values are expressed in mN/mm^2^.

| Twitch Contraction (mN/mm<sup>2</sup>) |  |  |  |
| --- | --- | --- | --- |
| Muscle | Resting Tension | Peak-Tension | Active Tension |
| EDL | $4.84 \pm 0.57$ | $24.24 \pm 2.15$ | $19.4 \pm 2.72$ |
| SOL | $12.08 \pm 0.93$ | $25.08 \pm 2.36$ | $13 \pm 3.29$ |
| Tetanic Contraction (mN/mm<sup>2</sup>) |  |  |  |
| Muscle | Resting Tension | Plateau Tension | Active Tension |
| EDL | $4.61 \pm 0.57$ | $89.04 \pm 8.37$ | $84.43 \pm 8.94$ |
| SOL | $10.64 \pm 1.49$ | $104 \pm 8.08$ | $93.36 \pm 9.57$ |

**Figure 4.**
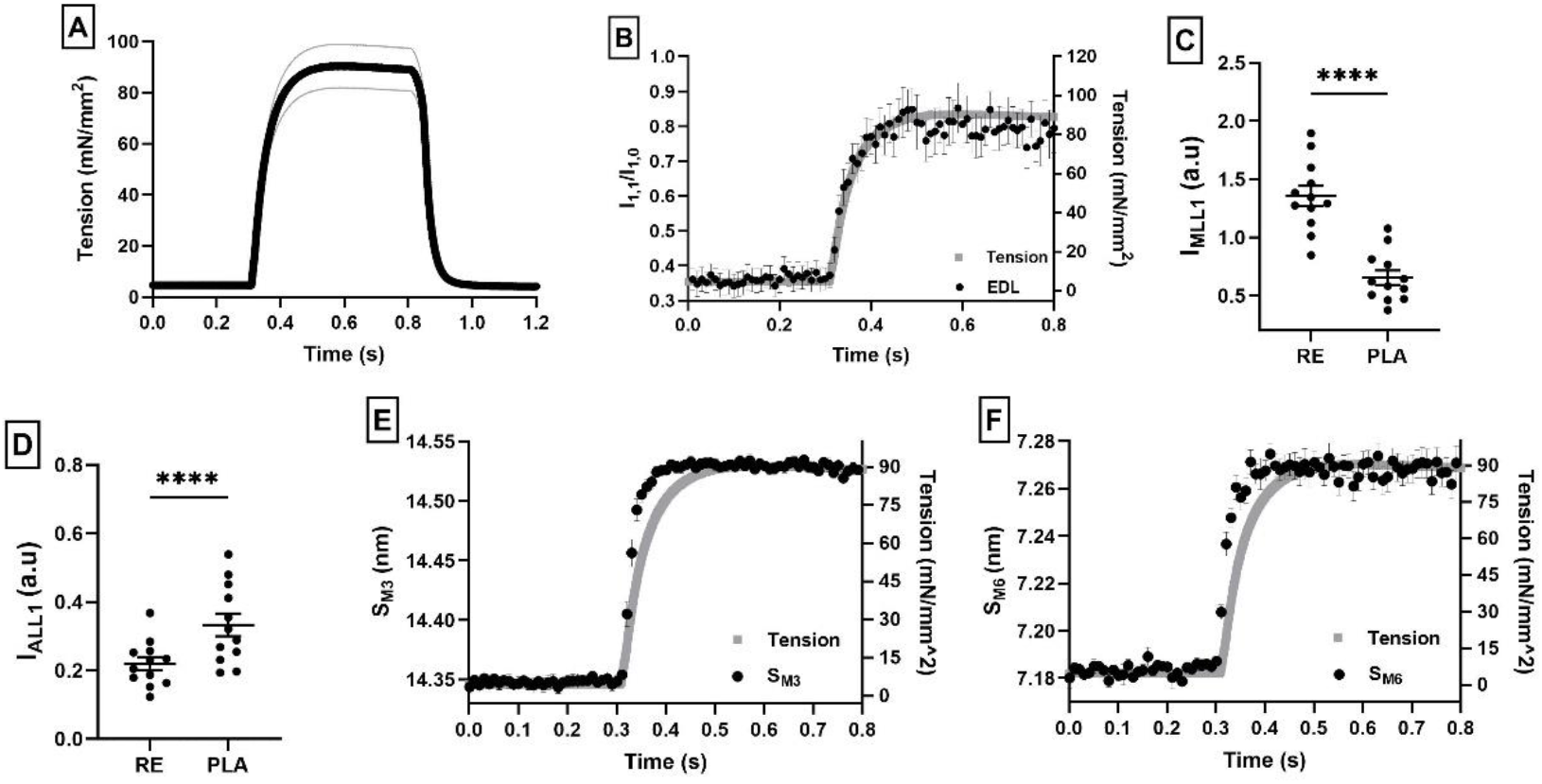
X-ray reflections of EDL muscle during tetanic contraction. As tension (A) develops, the intensity ratio (B), the spacing of M3 (S_M3_, E), and the spacing of M6 (S_M6_, F) all increase. The corresponding MLL1 intensity (I_MLL1_, C) decreases and ALL1 intensity (I_ALL1_, D) increases. Stimulation frequency was 100 Hz. All data are plotted as mean ± SEM, n = 8.

**Figure 5.**
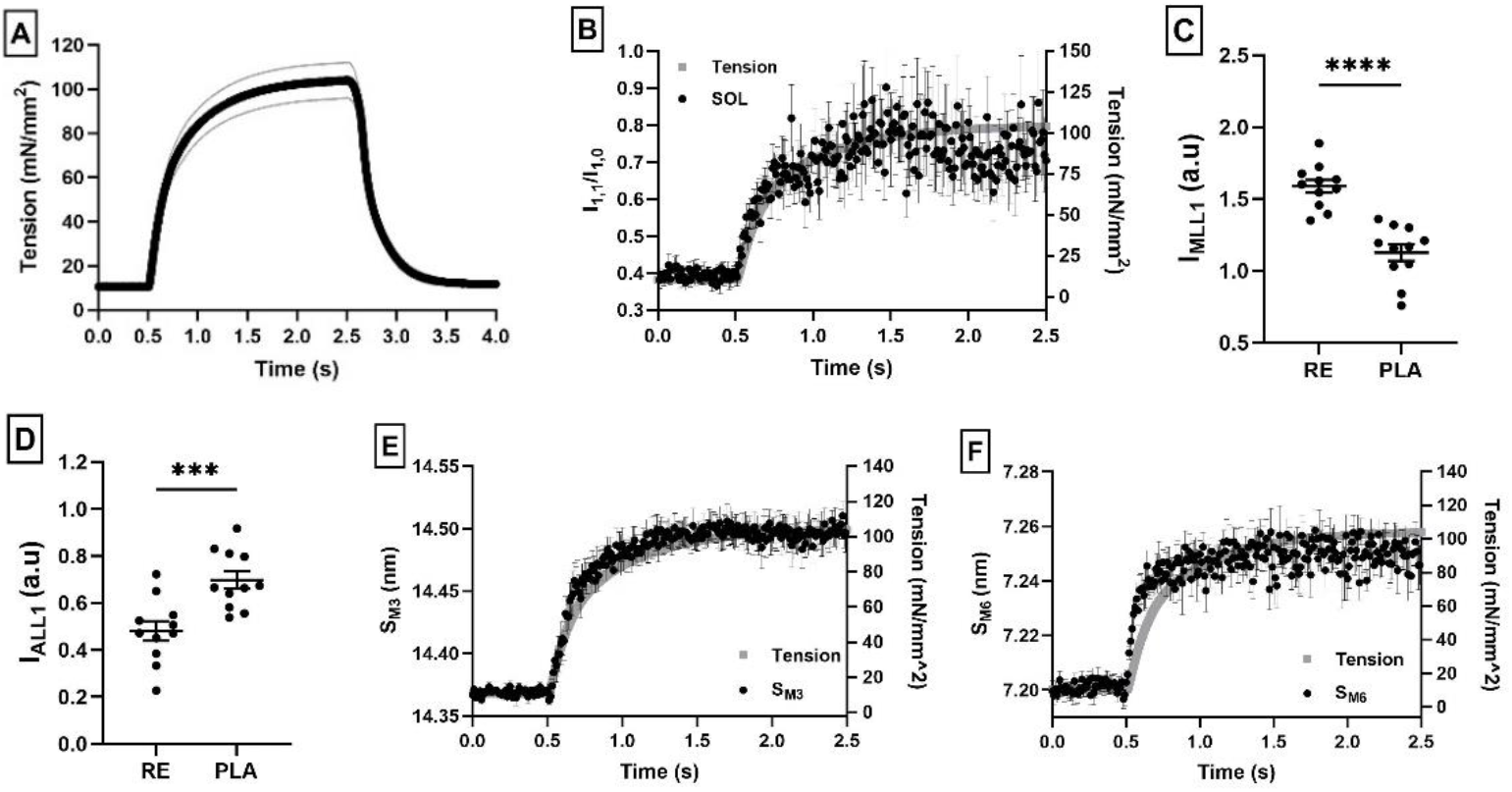
x-ray reflections of SOL during tetanic contraction. As tension (A) develops, the intensity ratio (B), the spacing of M3 (S_M3_, E), and the spacing of M6 (S_M6_, F) all increase. The corresponding MLL1 intensity (I_MLL1_, C) decreases and ALL1 intensity (I_ALL1_, D) increases. Stimulation frequency was 100 Hz. All data are plotted as mean ± SEM, n = 7.

The intensity ratio (I_1,1_/I_1,0_) from EDL muscle was initially 0.36 ± 0.03 at rest, increasing with a half time of 35.8 ms (95% CI: 28 ms to 44 ms) to 0.83 ± 0.05 at the tension plateau (p < 0.0001, Figure 4B). For SOL muscle, I_1,1_/I_1,0_ increased from 0.38 ± 0.01 at rest to 0.74 ± 0.06 at the tension plateau (p = 0.0003, Figure 5B) with a half-time of 121.6 ms (95% CI: 87.7 ms to 159.6 ms). Comparing to twitch, the change in I_1,1_/I_1,0_ during tetanus is 3.6 times more for EDL and 4.5 times more for SOL. The rate of tension development in EDL is 21.6% slower than the increase in I_1,1_/I_1,0_, and 38.3% slower for SOL muscle. This observation agrees with previous reports, suggesting the myosin heads in EDL and SOL move towards the thin filament ahead of force production (Huxley, 1968; Haselgrove & Huxley, 1973; Cecchi *et al*., 1991; Hill *et al*., 2021).

I_MLL1_ from EDL muscle decreased significantly from 1.36 ± 0.09 at rest to 0.65 ± 0.06 at maximum contraction (p < 0.0001, Figure 4C) whereas I_ALL1_ increased from 0.22±0.02 to 0.33±0.03 (P < 0.0001, Figure 4D). I_MLL1_ from SOL muscle decreased from 1.59 ± 0.05 at rest to 1.13 ± 0.06 at the tension plateau (p < 0.0001, Figure 5C) and I_ALL1_ increased from 0.48 ± 0.04 to 0.70 ± 0.04 (p < 0.0006, Figure 5D). These results suggest that, in both SOL and EDL muscle, the myosin heads depart from the region of the thick filament backbone and interact with the thin filament to form crossbridges consistent with previous reports (Reconditi *et al*., 2011; Caremani *et al*., 2019; Hill *et al*., 2021).

During tetanic contraction, the axial periodicity of the myosin heads (∼14.3 nm at rest) increased ∼1% to ∼14.5 nm. This has been interpreted as the myosin heads transitioning from a resting (OFF) to its active (ON) configuration able to produce force (Linari et al. 2015). In EDL muscle, S_M3_ increased ∼1.2% from 14.35 ± 0.005 nm to 14.53 ± 0.004 nm at maximum tension (p < 0.0001, Figure 4E). For SOL muscle S_M3_ increased ∼0.8% from 14.37 ± 0.001 nm at rest to 14.49 ± 0.01 nm at maximum tension (p < 0.0001, Figure 5E. The half-time of the change in S_M3_ from rest to plateau was 27.6 ms (95% CI: 26.1 ms to 29.1 ms) and 139.3 ms (95% CI: 125.4 ms 153.8 ms) for EDL and SOL, respectively. Compared to the half-times of tension development, the rise of tension is 39% slower than the rise of S_M3_ in EDL and 29% in SOL (Piazzesi *et al*., 1999; Reconditi *et al*., 2011; Ma *et al*., 2018*b*).

S_M6_ changed from 7.19 ± 0.002 nm to 7.27 ± 0.005 nm (p < 0.0001, Figure 4F) at maximum contraction for EDL. In SOL, S_M6_ increased from 7.20 ± 0.003 nm at rest to 7.25 ± 0.005 nm (p < 0.0001, Figure 5F). The backbones of the thick filaments in both muscles are extended to a greater extent during tetanic contraction than during twitch contraction, consistent with the greater active force developed during tetani. The half-time of S_M6_ development is 16.3 ms (95% CI: 13.4 ms to 19.4 ms) for EDL, and 53.8 ms (95% CI: 39.6 ms to 69.6 ms) for SOL. Comparing the half-time of tension development to the rise of S_M6_, the rise of tension is 64% slower in EDL, and 73% slower in SOL (Reconditi *et al*., 2011; Ma *et al*., 2018*b*; Hill *et al*., 2021).

### Changes in thick filament structure as a function of tension

The mechano-sensing thick filament activation model posits a distinct structural change in the thick filament at a particular level of active tension (Linari *et al*., 2015; Irving, 2017; Ma *et al*., 2018*b*). We examined this notion by examining the changes of S_M3_ (ΔS_M3_) and S_M6_ (ΔS_M6_) during the rising phase of tetanic contraction as a function of active tension (Figure 9). The rising phase is defined from the start of the stimulation to the inflection point where the tension starts to plateau.

For EDL the rising phase is defined from 0.3 s to 0.47 s and 0.5 s to 1.4 s for SOL muscle. When plotting ΔS_M6_ of EDL and SOL as a function of active tension, both traces have a non-linear relationship, that may be fit with a Hill function, with active tension (Figure 6A and B, opened gray squares). This finding agrees with previous observations made in mouse (Ma *et al*., 2018*b*) and frog (Linari et al. 2015; Brunello et al. 2014) skeletal muscle.

**Figure 6.**
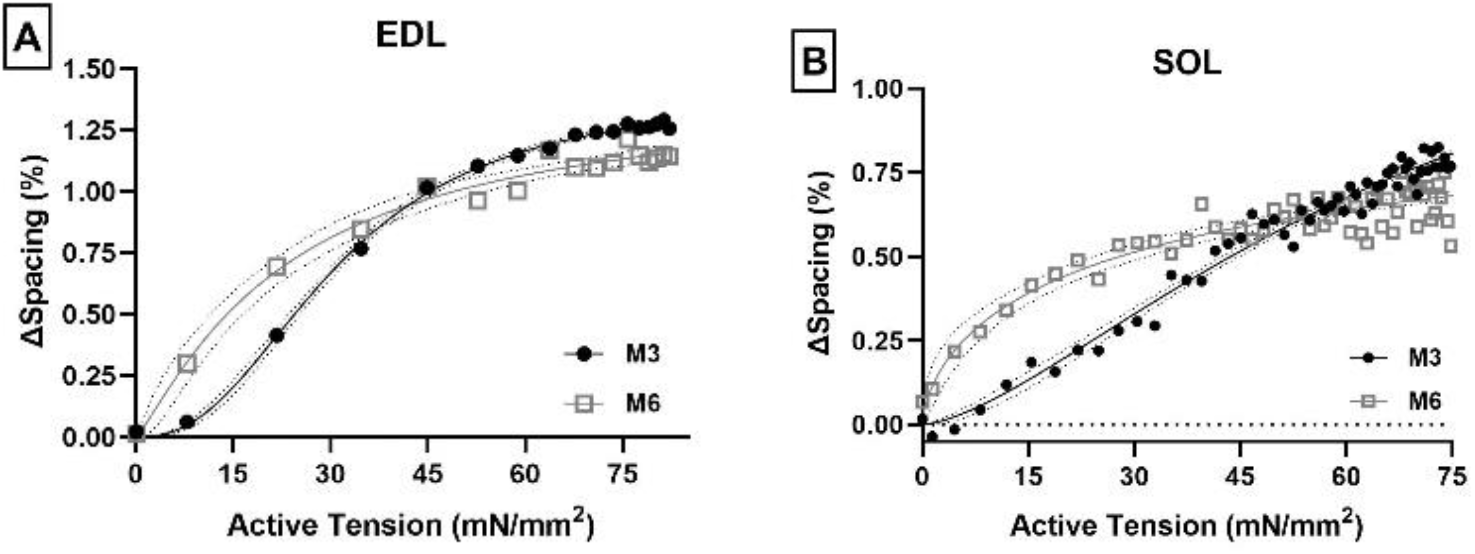
ΔS_M3_ and ΔS_M6_ plotted against active tension of EDL and SOL muscle. The ΔS_M3_ (opened gray square) and ΔS_M6_ (filled black circles) in EDL (A) muscle have a non-linear relationship with active tension. The ΔS_M6_ of SOL (B) muscle also has a similar non-linear shape (opened gray squares), but ΔS_M3_ appears much less sigmoidal (filled black circles).

ΔS_M3_ of EDL muscle as a function of active tension (Figure 6A, filled black circles) appears to have a sigmoidal shape similar to that reported previously for mouse EDL muscle (Ma *et al*., 2018*b*). This curve is well fit by a Hill function (R^2^ = 0.9988) with a Hill coefficient of 2.424 indicative of a cooperative process. In contrast, the ΔS_M3_ of SOL muscle as a function of active tension (Figure 6B, filled black circles) appears to be less cooperative with a significantly smaller Hill coefficient (h = 1.522, p=0.0004) than in EDL, so that the relation appears close to linear with no indication of saturation at high tension, indicating that the changes in the disposition of myosin heads with tension are much less cooperative in SOL. These observations suggest that the myosin heads in SOL muscle may be regulated differently than EDL muscle.

ΔS_M6_ has a non-linear relationship with active tension in both EDL (Figure 6A, open gray squares) and SOL (Figure 6B, open gray squares) indicating the extensibilities of the backbone of the thick filament are a non-linear function of tension in both muscles similar to previous observations in mouse (Ma *et al*., 2018*b*). The maximum extensibility of SOL muscle, however, is much less than in EDL for the same amount of active tension suggesting that thick filaments are stiffer in SOL than in EDL. To resolve this question, we estimated thin filament extensibilities in both SOL and EDL muscles. Given that the thin filament extensibility is a linear function of active force (Kiss *et al*., 2018), changes in the 2.73 nm reflection spacing can be used as a measure of the local force experienced by the individual myofilaments in the sarcomeres (Mijailovich *et al*., 2019). To estimate the thin filament extensibility, the axial spacing of actin subunits was estimated from the S_ALL6_ and S_ALL7_ values (see methods). The axial displacement of EDL increased 0.39% (2.731 ± 0.001 nm to 2.742 ± 0.001 nm, p < 0.0001, Figure 7A) and increased 0.30% (2.733 ± 0.001 nm to 2.741 ± 0.001 nm, p < 0.0001, Figure 7B) in SOL muscle (Wakabayashi *et al*., 1994; Kiss *et al*., 2018). The changes in spacing are not significantly different in EDL and SOL (p = 0.1612, Figure 7C) and are comparable with previous reported values from frog muscle and mice (Wakabayashi *et al*., 1994; Kiss *et al*., 2018). This observation indicates that the extensibilities of thin filaments in EDL and SOL are not significantly different during tetanic contraction suggesting that the strain generated in the thin filaments due to force development by the myosin heads was similar in both muscles and that the force experienced by the individual myofilaments in the sarcomeres is also similar. In contrast to the change in actin periodicities, the change of thick filament backbone periodicity during contraction is significantly different (p = 0.0002, Figure 7D) between EDL (1.17%) and SOL (0.7%). This suggests that the results reported in Figure 6 reflect differences in the intrinsic properties (stiffness) of the thick filaments in the two muscles.

**Figure 7.**
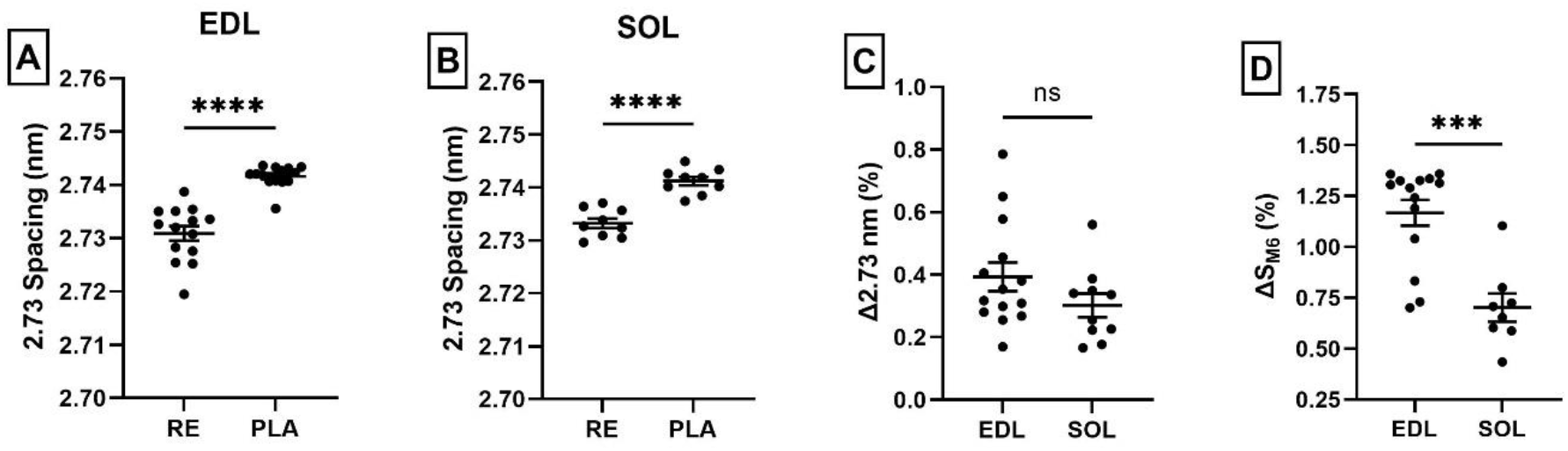
Axial rise of actin monomers of EDL and SOL during tetanic contraction. The axial spacing of two actin monomers (2.73 nm) in EDL (A) and SOL (B) increased from resting to the plateau region. The change of 2.73 nm reflection is not significant between EDL and SOL (C). On the other hand, the change of the backbone of the thick filament was significant difference between EDL and SOL (D). Error bars indicate mean ± SEM.

## Discussion

### Overview

A mechano-sensing mechanism for thick filament activation has been proposed based on observations made in frog fast skeletal muscle (Linari et al. 2015). These authors argued that strain in the thick filament backbone is the molecular switch that activates the thick filament. Since then, evidence for this mechano-sensing mechanism has been obtained from rat cardiac muscle (Caremani *et al*., 2018), and mouse EDL skeletal muscle (Hill *et al*., 2021). Studies of the applicability of this model to slow skeletal muscle have been lacking. Here we investigated the structural changes of the thick filament associated with thick filament regulatory processes comparing predominately slow-twitch SOL and pure fast-twitch EDL from the rat. The results we obtained imply that thick filament activation mechanisms proceed differently in the two muscle types.

### Structural events during twitch contractions in fast and slow rat skeletal muscle

The spacing of the 6^th^ order myosin based meridional reflection, S_M6_, from EDL muscle increases significantly during rise of tension during a twitch contraction followed by reductions in I_MLL1_ and increase in S_M3_ consistent with the mechano-sensing model (Figure 2). During twitch contractions in SOL muscle, however, while S_M6_ increased with tension similar to in EDL muscle (Figure 3F), S_M3_ remained unchanged as tension rises (Figure 3E). The lack of change in S_M3_ suggests that the majority of the myosin heads do not transition from their OFF states toward the ON state in response to twitch stimulation. The lack of change in I_MLL1_ also agree with this interpretation, indicating that the degree of helical ordering of the myosin heads on the thick filament is preserved at peak-tension (Figure 3C and D). However, there must be some degree of cross-bridge formation to generate the observed force and the increased association of myosin heads with the thin filament as indicated by the small rise in I_1,1_/I_1,0_ (Figure 3A and B). The population of myosin heads that contribute to the production of force might consist of constitutively ON heads (Irving, 2017) in SOL muscle that responded to the twitch signal.

### Structural events during tetanic contractions fast and in slow rat skeletal muscle

We have shown that EDL muscle exhibits significant alterations in myosin head configuration during twitch contractions, whereas myosin heads in SOL muscle retained most of their quasi-helical arrangement around the thick filament backbone throughout the twitch. Our results from tetanic contractions provide further insights into the molecular basis of this different behavior.

The X-ray diffraction results from rat EDL muscle during tetanic contraction, as shown above for twitches, follow the expected sequence of events from the mechano-sensing mechanism. During tetanic contraction, the periodicity of the backbone of the thick filament (S_M6_) increased ∼1.1% from its resting state (Figure 4F) as seen in many previous studies of skeletal muscle (Huxley & Brown, 1967; Haselgrove, 1975; Huxley *et al*., 1994; Wakabayashi *et al*., 1994; Linari *et al*., 2015). These changes in S_M6_ appear to lead the rise in tension, but it could reflect a nonlinear response of S_M6_ to tension (Ma et al., 2018). The changes in S_M6_ are followed by an increase in the axial spacing (∼1.2%) of the crowns of myosin heads (S_M3_, Figure 4E) as a result of crossbridge formation leading to the rise of tension indicating that most myosin heads are in the ON state (Huxley & Brown, 1967; Huxley, 1973; Linari *et al*., 2015). The transition of S_M3_ to ON state is a cooperative transition as demonstrated by the Hill fits in Figure 6A. This cooperative change in S_M3_ is accompanied by a reduction in the helical ordering of the myosin heads on the thick filament (I_MLL1_, Figure 4C) at the plateau region of the tension curve, an increase in the intensity of actin-based layer line (I_ALL1_, Figure 4C), and an increase in I_1,1_/I_1,0_ (Figure 4B) (Huxley, 1973; Haselgrove & Huxley, 1973; Haselgrove, 1975; Bordas *et al*., 1999; Tamura *et al*., 2009). These results are consistent with recent studies of mouse EDL muscle (Hill *et al*., 2021).

In tetanic contraction of SOL muscle, unlike twitch contractions, the changes in x-ray reflections between rest to the plateau of isometric tension are also consistent with an OFF to ON transition in the thick filaments but are blunted in comparison to EDL muscle. S_M3_ in SOL muscle increased ∼0.8%, as compared to 1.2% in EDL from their resting values (Figure 5E). Also unlike twitches, there is a significant reduction in I_MLL1_ by ∼ 39%, as compared to ∼52% in EDL indicating disruption of the helical arrangement of the myosin heads at the plateau region, indicating a substantial population of myosin heads are no longer tightly bound to the thick filament in the ordered state (Figure 5C) (Haselgrove, 1975) and are now in an active configuration (Huxley & Brown, 1967; Huxley, 1973; Linari *et al*., 2015). The population of myosin heads that is no longer in the ordered state has left the thick filament and are in the vicinity of the thin filament as indicated by the significant increase in I_1,1_/I_1,0_ (Figure 5B), by 195%, as compared to 230% in EDL. All of these changes in the diffraction features associated with OFF to ON transitions in the thick filaments are less in SOL than in EDL muscle. The most dramatic difference, however, between the behavior of the X-ray reflections in EDL and SOL muscle during tetanus is that the transition of S_M3_ to active configuration in SOL muscle is less cooperative as indicated by the reduced Hill coefficient and the lack of a plateau at high tension so that the relationship is more linear. Together, these results indicate significant differences in thick filament regulation in SOL as compared to EDL muscle.

### Implications for thick filament activation

The differences in thick filament activation in EDL and SOL muscle could be due to a number of factors including differences in the helical packing of the myosin heads along the thick filament leading to differences in the strength of head-backbone (intermolecular) and head-head (intramolecular) interactions (Craig & Padrón, 2022). Another possibility is the differences in backbone structure. ΔS_M6_ in both EDL and SOL muscle is a non-linear function of the increase in tension (Figure 6) (present study; Ma et al. 2018). The maximum extension of thick filaments in SOL muscle, however, is much less than in EDL at the same level of developed force at the myofilament level (same degree of thin filament extension). The simplest explanation for this observation is that thick filaments in SOL muscles are substantially stiffer than in EDL. Backbone elongation is assumed to be the molecular trigger in the mechano-sensing mechanism. A stiffer thick filament backbone in SOL muscle could provide a molecular explanation for why SOL lacks evidence for mechano-sensing during twitch, and the absence of cooperative transitions in myosin head periodicity in tetani, if the thick filament extension in SOL is insufficient to trigger such cooperative transitions.

There may be an alternative mechanism in terms of the direct response of thick filaments to calcium in fast and slow skeletal muscle. We have recently demonstrated in porcine left ventricular myocardium (Ma *et al*., 2022) that cooperative OFF/ON transitions in the thick filament can be induced directly by calcium in the absence of thick filament strain. As calcium levels rise during contraction, myosin heads will be released from the thick filament backbone at the same time that regulatory units open up on the thin filament. If there were differences in the thick filament response to calcium between SOL and EDL muscle, this could help explain the differences we see in thick filament activation. Such differences may be revealed in future studies.

## Conclusions

In this study we compared the dynamic changes in sarcomere structure in intact fast-twitch EDL muscle and slow-twitch SOL muscle during twitch and tetanic contraction. During twitch contraction, at optimal length, EDL muscle exhibits significant changes in S_M3_, S_M6_, and I_MLL1_ indicating a transition of myosin heads to an active ON configuration. In contrast, SOL muscle did not show significant changes in S_M3_ and I_MLL1_ as in EDL muscle indicating that the majority of myosin heads retain their quasi-helical ordering along the thick filament during twitch. In tetanic contraction, both SOL and EDL show significant changes in I_11_/I_10_, I_MLL1_, S_M6_ and S_M3_ demonstrating the thick filaments have been turned ON but the relative magnitude of these changes in less in SOL muscle. ΔS_M3_ in EDL has a sigmoidal relationship with active tension indicating that the transition of myosin heads from OFF to ON is a cooperative process as expected by the mechano-sensing mechanism. In contrast, ΔS_M3_ as a function of active tension in SOL appears more linear with less cooperativity as compared to EDL.

The extensibility of the backbone, as indicated by ΔS_M6,_ is a non-linear function of active tension in both muscles as observed previously for other vertebrate muscle (Linari *et al*., 2015; Kiss *et al*., 2018; Ma *et al*., 2018*b*). The trigger in the mechano-sensing model is proposed be the elongation the thick filament backbone in response to active force production. The extension of the backbone of the thick filament backbone at maximum tension in SOL, however, was significant less than in EDL due to intrinsic differences in the backbone of thick filament between SOL and EDL muscle that result in greater effective stiffness of the SOL thick filament. A stiffer thick filament backbone in SOL provides a molecular explanation for the lack of a classic mechano-sensing response during twitches. The lower cooperativity and lack of a plateau in the ΔS_M3_/force relationship during force development in tetani suggest that thick filament activation proceeds differently in slow muscle, with the OFF-ON transition being a continuous process, rather than a stepwise, cooperative process as in fast EDL muscle. In fast muscle, the ability to quickly turn the myosin heads ON in a stepwise, cooperative manner, is consistent with its physiological role of generate maximum power in a short period of time. Considering the important physiological role of slow skeletal muscle in maintaining posture, it would seem advantageous for the muscle to regulate its myosin heads in a more finely tuned, graded manner.

## Competing interests

T.I provides consulting and collaborative research studies to Edgewise Therapeutics and Bristol Myers Squibb, but such work is unrelated to the content of this article. Other authors declare no conflict of interests.

## Author Contributions

W.M, M.R., T.I designed the experiments; H.G, W.M performed the research; H.G analyzed the data; H.G, W.M, T.I, and M.R wrote the manuscript. All authors approved the final version of the manuscript.

## Funding

This research used resources of the Advanced Photon Source; a U.S. Department of Energy (DOE) Office of Science User Facility operated for the DOE Office of Science by Argonne National Laboratory under Contract No. DE-AC02-06CH11357. This project is supported by grant P30 GM138395 from the National Institute of General Medical Sciences of the National Institutes of Health. Funding for MR comes from NIH R01HL128368 and the UW Center for Translational Muscle Research, sponsored by NIH NIAMS P30AR074990.The content is solely the authors’ responsibility and does not necessarily reflect the official views of the National Institute of General Medical Sciences or the National Institutes of Health.

## Acknowledgements

We thank Vitold E. Galkin from Easter Virginia Medical School and Anthony L. Hessel from Westfalische Wilhelms-Universitat Munster for providing helpful comments to the manuscript.

