## Supplementary material for "Thick Filament Activation Is Different in Fast and Slow-Twitch Skeletal Muscle": SI

Table of Contents Category: Muscle

### Supplemental Results

#### Structural changes during fixed-end twitch contractions in EDL and Soleus muscle

The inter-filament lattice spacing, $d_{1,0}$, is measured from the distance between the equatorial 1,0 reflections to the center of the pattern. When the muscle was stimulated, $d_{1,0}$ increased from 34.01 ± 0.22 nm to 34.36 ± 0.24 nm ($p < 0.0001$, Figure S1A) for EDL muscle and increased from 34.36 ± 0.2 nm to 34.58 ± 0.23 nm ($p = 0.0021$, Figure S2A) in SOL muscle. The increase of $d_{1,0}$ from rest to peak-tension in both muscles suggest the lattice spacing has expanded during contraction as a result of internal shortening of the sarcomeres that is known to occur when the muscle as a whole is kept at fixed length (Julian et al., 1978; Kawakami & Lieber, 2000). Assuming constant sarcomere lattice volume throughout muscle contraction in intact preparations (Elliott et al., 1963; Millman, 1998), the change of sarcomere length (SL) can be estimated based on the changes in $d_{1,0}$. Using this assumption, the lattice spacing changes would imply a decrease in SL from 2.73 ± 0.03 µm at rest to 2.67 ± 0.03 µm (~2%) at peak-tension (Figure S1B) in EDL muscle and from 2.81 ± 0.03 µm at rest to 2.75 ± 0.04 µm (~2%) at peak-tension (Figure S2B) in SOL muscle. These estimated changes in SL are smaller compared to previous reported values from rat EDL and SOL muscles (Mutungi & Ranatunga, 2000). However, the shape of the SL curves, where SL remain shortened while tension declines, is similar to that observed previously (Mutungi & Ranatunga, 2000).

The intensity of the third order myosin meridional reflection ($I_{M3}$), after correction by multiplying by the radial width of the reflection (Huxley et al., 1982), increased from 1.74 ± 0.24 to 2.04 ± 0.17 at peak tension ($p = 0.1093$, Figure S1C) in EDL muscle whereas $I_{M3}$ in SOL muscle decreased from 1.75 ± 0.23 to 1.57 ± 0.16 at peak tension ($p = 0.2230$, Figure S2C). The intensity of the sixth order myosin meridional reflection ($I_{M6}$) did not change significantly during the twitch in both muscles (EDL 0.043 ± 0.007 rest to 0.045 ± 0.007 peak tension, $p = 0.7501$, Figure S1D and SOL 0.035 ± 0.008 to 0.038 ± 0.008, $p = 0.5358$, Figure S2D).

The thin filament is formed by two approximately helical strands of actin monomer intertwined with one another to form a double helical filament with a helical repeat of ~35.9 nm. The double helix can also be described as two genetic helices, a left-handed helix with a pitch of 5.9 nm generating the sixth actin-based layer line (ALL6), and a right-handed helix with a pitch of

5.1 nm, generating the seventh actin-based layer line (ALL7). The ALL7 is a weaker reflection than the ALL6 making it more difficult to measure accurately. The ALL6 and ALL7 were analyzed as described previously (Wakabayashi et al., 1994). For EDL, the spacing of the ALL6 reflection ($S_{ALL6}$) increased significantly from 5.88 ± 0.004 nm to 5.89 ± 0.003 nm (.17%) (p = 0.0018, Figure S1E), and its corresponding intensity ($I_{ALL6}$) also increased significantly from 0.48 ± 0.03 to 0.53 ± 0.02 (p = 0.0448, Figure S1F). The spacing of ALL7 ($S_{ALL7}$) did not change significantly going from 5.09 ± 0.005 nm to 5.092 ± 0.004 nm (p = 0.1284, Figure S1G). The intensity of ALL7 ($I_{ALL7}$) increased, but not significantly, from 0.19 ± 0.02 to 0.23 ± 0.03 (p = 0.2292, Figure S1H). In SOL muscle, $S_{ALL6}$ did not change significantly, going from 5.899 ± 0.03 nm to 5.893 ± 0.003 nm (p = 0.2482, Figure S2E), and the corresponding intensity did not change significantly going from 0.5 ± 0.06 to 0.53 ± 0.01 (p = 0.6059, Figure S2F). Neither $S_{ALL7}$ nor $I_{ALL7}$ changed significantly with $S_{ALL7}$ going from 5.09 ± 0.005 nm to 5.1 ± 0.002 nm (p = 0.0901, Figure S2G) and $I_{ALL7}$ going from 0.26 ± 0.03 to 0.29 to 0.03 (p = 0.5097, Figure S2H). The lack of significant changes in $S_{ALL6}$ and $S_{ALL7}$ reflections in SOL muscle indicate that the structure of the thin filament did not change appreciably during twitch. In EDL muscle, the significant change in $S_{ALL6}$ but not in $S_{ALL7}$ suggest the right-hand helix and the left-hand helix was extended to different amounts during twitch contraction indicating slight twisting of the thin filament during contraction (Wakabayashi et al., 1994).

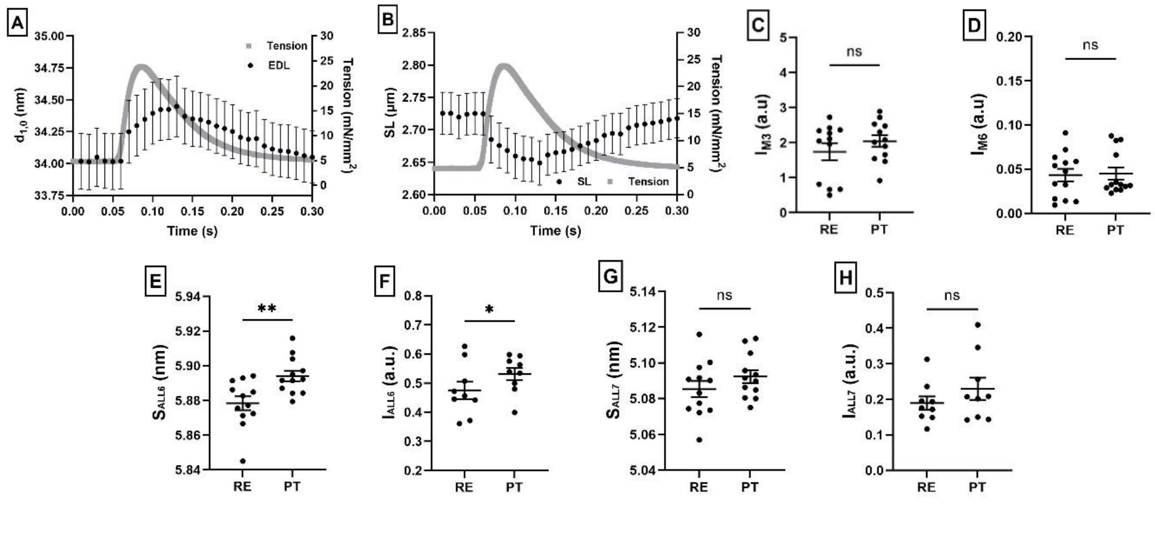


**Figure S1. EDL muscle during twitch contraction**. A. Changes in $d_{1,0}$ inter-filament lattice spacing plotted along with tension as a function of time during contraction. B. Estimated changes of sarcomere length (SL) plotted along with tension as a function of time during contraction. C. Change of the M3 intensity ($I_{M3}$) between rest (RE) and peak-tension (PT). D. Change of the M6 intensity ($I_{M6}$) between rest (RE) and peak-tension (PT). E. Spacing change of the sixth actin-based layer line ($S_{ALL6}$) between rest (RE) and peak-tension (PT) F. Change of ALL6 intensity ($I_{ALL6}$) between rest (RE) and peak-tension (PT). G. Spacing change of the seventh actin-based layer line ($S_{ALL7}$) between rest (RE) and peak-tension (PT). H. Change of ALL7 intensity ($I_{ALL7}$) between rest (RE) and peak-tension (PT).

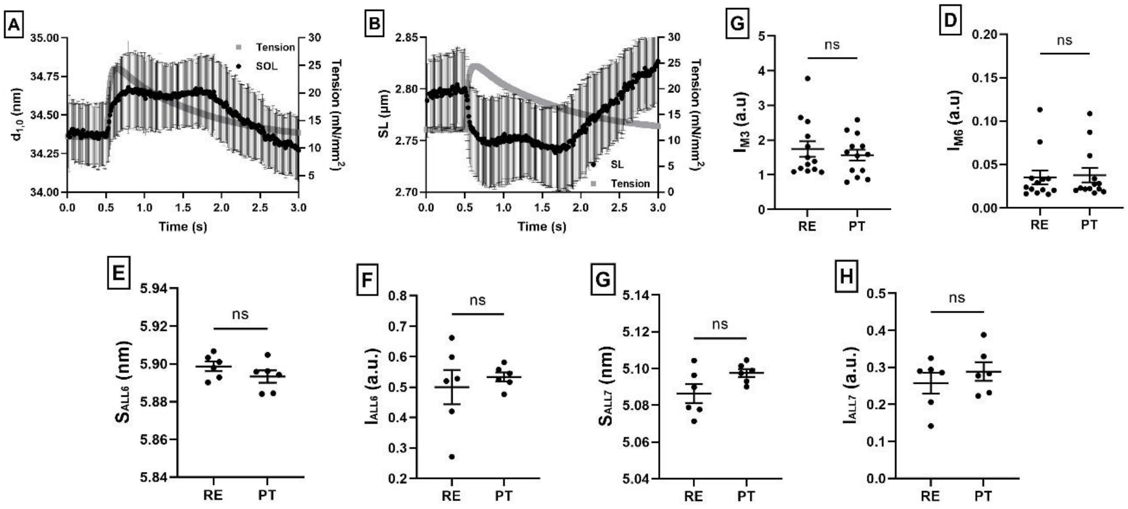


**Figure S2. SOL muscle during twitch contraction**. A. Change of $d_{1,0}$ inter-filament lattice spacing and developed tension plotted as a function of time during contraction. B. Estimated changes of sarcomere length (SL) and developed tension plotted as a function of time during contraction. C. Change of M3 intensity ($I_{M3}$) between rest (RE) and peak-tension (PT). D. Change of M6 intensity ($I_{M6}$) between rest (RE) and peak-tension (PT). E. Spacing change of the sixth actin-based layer line ($S_{ALL6}$) between rest (RE) and peak tension (PT). F. The intensity changes of the ALL6 ($I_{ALL6}$) between rest (RE) and peak tension (PT). G. Spacing change of the seventh actin-based layer line ($S_{ALL7}$) between rest (RE) and peak tension (PT) H. Intensity change of the ALL7 ($I_{ALL7}$) between rest (RE) and peak tension (PT)

### Structural changes during fixed-end tetanic contractions in EDL and Soleus muscle

When muscle undergo maximum tetanic contraction, the majority of the X-ray reflections changed more dramatically than in twitch contraction. The $d_{1,0}$ interfilament lattice spacing changed from 34.08 ± 0.19 nm to 35.18 ± 0.42 nm ($p = 0.0012$, Figure S3A) for EDL muscle, and 34.53 ± 0.23 nm to 35.98 ± 0.43 nm ($p = 0.0248$, Figure S4A) for SOL muscle. This decrease in $d_{10}$ during maximum tetanic contraction, as with twitch contractions, is consistent with some degree of sarcomere shortening under fixed end conditions. Assuming constant lattice volume, the sarcomere length of EDL is estimated to have decreased from 2.73 ± 0.03 µm at rest to 2.56 ± 0.05 µm (~6%) at maximum tension (Figure S3B). This change in SL in EDL is comparable to the value reported previously on rat EDL muscle (Mutungi & Ranatunga, 2000; Caremani et al., 2019). Similarly, sarcomere length in SOL muscle is estimated to decrease from 2.81 ± 0.03 µm at rest to 2.57 ± 0.05 µm (~9%) at maximum tension (Figure S4B) and this is greater than the value previously reported for rat SOL muscle (Mutungi & Ranatunga, 2000). However, the change in SL in both muscles are smaller compared to tibialis anterior muscle in mice (Moo & Herzog, 2020).

The intensity of the M3 myosin meridional reflection ($I_{M3}$) from EDL muscle, after width correction, increased significantly from 17.45 ± 1.84 at rest to 34.55 ± 2.75 at maximum tension ($p = 0.0017$, Figure S3C). $I_{M3}$ from SOL muscle increased from 13.41 ± 1.61 at rest to 20.06 ± 2.59 at maximum tension ($p = 0.0395$, Figure S4C). The intensity of the M6 myosin meridional reflection ($I_{M6}$) decreased, but not significantly, from 0.29 ± 0.04 at rest to 0.15 ± 0.01 at maximum tension ($p = 0.5358$, Figure S3D) for EDL muscle and 0.18 ± 0.015 at rest to 0.11 ± 0.003 at maximum tension ($p = 0.0010$, Figure S4D) for SOL muscle. The significant changes in $I_{M3}$ and $I_{M6}$ in both muscles indicate there were substantial structural changes in thick filament structure going from the rest to maximum contraction.

The ALL6 reflection from the left-handed actin helix showed a significant increase in spacing from 5.89 ± 0.004 nm to 5.91 ± 0.002 nm (0.34%) ($p < 0.0001$, Figure S3E) in EDL muscle. The intensity of the ALL6 actin reflection ($I_{ALL6}$) also increased for EDL muscle from 0.4 ± 0.02 to 0.6 ± 0.04 ($p < 0.0001$, Figure S3F). The spacing of the ALL7 reflection from the right-handed actin helix increased from 5.09 ± 0.005 nm at rest to 5.11 ± 0.002 nm (0.39%) at maximum tension ($p < 0.0001$, Figure S3G) and its intensity increased from 0.13 ± 0.01 at rest to 0.19 ± 0.02 at maximum tension ($p < 0.0001$, Figure S3H). For SOL muscle, $S_{ALL6}$ changed from 5.89 ± 0.002 nm to 5.91 ± 0.001 nm (0.34%) ($p < 0.0001$, Figure S4E). $I_{ALL6}$ increased from 0.37 ± 0.02 to 0.47 ± 0.03 ($p = 0.006$, Figure S4F). $S_{ALL7}$ increased from 5.097 ± 0.002 nm to 5.113 ± 0.002 nm (0.26 %) ($p < 0.0001$, Figure S4G). Lastly, $I_{ALL7}$ of SOL changed from 0.12 ± 0.01 to 0.19 ± 0.01 ($p = 0.0118$, Figure S4H). The significant increases of $I_{ALL6}$ and $I_{ALL7}$ in both muscles are consistent with cross-bridge formation adding to the thin filament mass in the form of bound myosin heads (Tsaturyan et al., 2011). $S_{ALL6}$ and $S_{ALL7}$ was extended to a longer length at plateau compare to resting indicating that force generated by the myosin heads causes slight extension of the thin filaments during force production (Huxley et al., 1994; Wakabayashi et al., 1994).

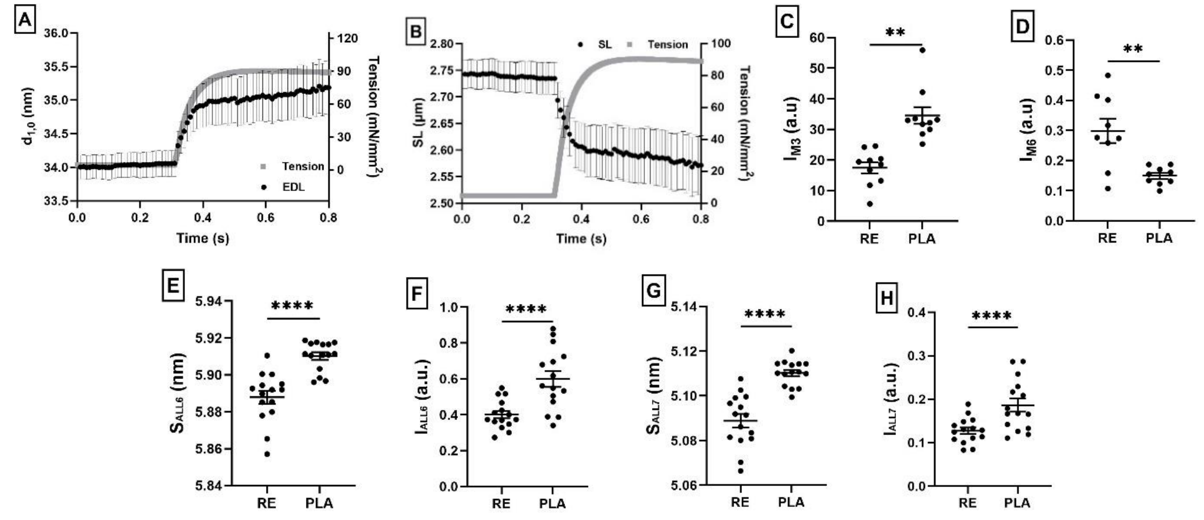


**Figure S3. EDL muscle during tetanic contraction**. A. The changes of $d_{1,0}$ inter-filament lattice spacing plotted along with tension as a function of time during contraction. B. Estimated changes of sarcomere length (SL) plotted along with tension as a function of time during contraction. C. Change of M3 intensity ($I_{M3}$) between rest (RE) and the tension plateau (PLA)n. D. Change of M6 intensity ($I_{M6}$) between rest (RE) and the tension plateau (PLA). E. The spacing change of sixth actin-based layer line ($S_{ALL6}$) between rest (RE) and the tension plateau (PLA). F. Intensity changes of ALL6 ($I_{ALL6}$) between rest (RE) and the tension plateau (PLA). G. Spacing change of the seventh actin-based layer line ($S_{ALL7}$ between rest (RE) and the tension plateau (PLA). H. Intensity changes of ALL7 ($I_{ALL7}$) between rest (RE) and the tension plateau (PLA).

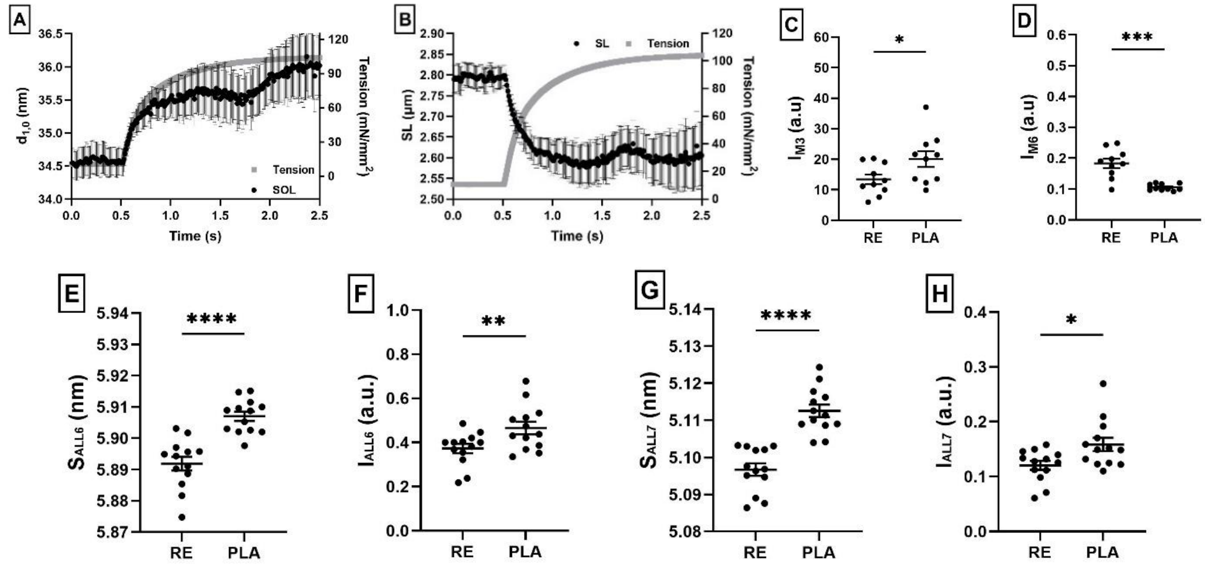


**Figure S4. SOL muscle during tetanic contraction**. Changes in $d_{1,0}$ inter-filament lattice spacing plotted along with tension during contraction. B. Estimated changes of sarcomere length (SL) plotted along with tension during contraction. C. Chnage of M3 intensity ($I_{M3}$) between rest (RE) and the tension plateau (PLA). D. Change of M6 intensity ($I_{M6}$) between rest (RE) and the tension plateau (PLA). E. Spacing change of the sixth actin-based layer line ($S_{ALL6}$) between rest (RE) and the tension plateau (PLA). F. Intensity changes of ALL6 ($I_{ALL6}$) een rest (RE) and the tension plateau (PLA). G. Spacing change of seventh actin-based layer line ($S_{ALL7}$) between rest (RE) and the tension plateau (PLA). H. Changes of ALL7 intensity ($I_{ALL7}$) between rest (RE) and the tension plateau (PLA).
